# Lower slow wave sleep and rapid eye-movement sleep are associated with brain atrophy of AD-vulnerable regions

**DOI:** 10.1101/2025.01.12.632386

**Authors:** Gawon Cho, Adam P. Mecca, Orfeu M. Buxton, Xiao Liu, Brienne Miner

**Affiliations:** Yale School of Medicine, New Haven, CT, U.S.A.; Yale Alzheimer’s Disease Research Unit, New Haven, CT, U.S.A.; The Pennsylvania State University, University Park, PA, U.S.A.

**Keywords:** sleep architecture, brain atrophy, inferior parietal region, Alzheimer’s disease, cerebral microbleeds

## Abstract

**Study objectives:** Sleep deficiency is associated with Alzheimer’s disease (AD) pathogenesis. We examined the association of sleep architecture with anatomical features observed in AD: (1) atrophy of hippocampus, entorhinal, inferior parietal, parahippocampal, precuneus, and cuneus regions (“AD-vulnerable regions”) and (2) cerebral microbleeds.

**Methods:** In 271 participants of the Atherosclerosis Risk in the Communities Study, we examined the association of baseline sleep architecture with anatomical features identified on brain MRI 13∼17 years later. Sleep architecture was quantified as the proportion of slow wave sleep (SWS), proportion of rapid eye-movement sleep (REM), and arousals index using polysomnography. Outcomes included (1) volumetric measurements of each AD-vulnerable region and (2) the presence of any cerebral microbleeds (CMBs) and that of lobar CMBs, which are more specifically associated with AD. We analyzed the association of each sleep predictor with each MRI outcome, adjusting for covariates.

**Results:** Having less SWS was associated with smaller inferior parietal region (β=−44.19 mm^3^ [95%CI=−76.63,−11.76]) and cuneus (β=−11.99 mm^3^ [−20.93,−3.04]) after covariate adjustment. Having less REM was associated with smaller inferior parietal region (β=−75.52 mm^3^ [−129.34, −21.70]) and precuneus (β=−31.93 mm^3^ [−63.79,−0.07]). After FDR adjustments, lower SWS and REM, respectively, were associated with smaller inferior parietal region. Arousal index was not associated with the volumes of AD-vulnerable regions. None of the sleep architecture variables were associated with CMBs or lobar CMBs.

**Conclusions:** Sleep deficiency is associated with the atrophy of the inferior parietal region, which is observed in early AD. Sleep architecture may be a modifiable risk factor for AD.

**Brief summary:** *Current Knowledge/Study Rationale: two sentences summarizing why the study was done:* While impaired sleep architecture has been associated with Alzheimer’s disease [AD] diagnosis and cognitive decline. To better understand the impact of sleep on AD pathogenesis, this study examined the association of sleep architecture with anatomical features observed in AD, including the atrophy of AD-vulnerable regions and CMBs.

*Study Impact: two sentences summarizing how the study impacts the field:* Our study shows that lower slow wave sleep and rapid eye movement sleep may be precipitating factors of inferior parietal region atrophy, which is associated with AD risk. Importantly, the current study’s findings can help characterize underlying mechanisms of how sleep deficiency, a prevalent disturbance among middle-aged and older adults, may facilitate AD pathogenesis and cognitive impairment.

## INTRODUCTION

Alzheimer’s disease (AD) is a progressive condition affecting over 6 million U.S. older adults. ^1^ It is characterized by distinct anatomical features, one of which is brain atrophy, a structural correlate of neurodegeneration. ^2–5^ During the preclinical and clinical phases of AD, brain regions have been observed to show different degrees of atrophy, which is often pronounced in the hippocampus, parahippocampal, entorhinal, inferior parietal, precuneus and cuneus regions (henceforth referred to as AD-vulnerable regions; see Figures 1a and 1b). ^2–6^ Prior research has shown that brain atrophy specific to these regions has been associated with increased conversion to AD and related dementias (from normal cognition and mild cognitive impairment), ^6–8^ as well as having severe neurocognitive impairment in AD. ^3,5^ Moreover, cerebral microbleeds (CMBs), which signify the presence of cerebral small vessel disease, ^9^ are another anatomical feature associated with AD (see Figure 1). ^10^ The presence of CMBs has been shown to predict subsequent cognitive impairment and the onset of AD and related dementias. ^11–16^ Among various types of CMBs, lobar microbleeds were most frequently observed in preclinical and early AD. ^15–17^ Lobar microbleeds have also been robustly associated with cortical amyloid-β load, a hallmark of AD pathology. ^15–17^ Identifying precipitating factors of these anatomical correlates of AD is necessary to gain a better understanding of potential modifiable risk factors.

**Figure 1.**
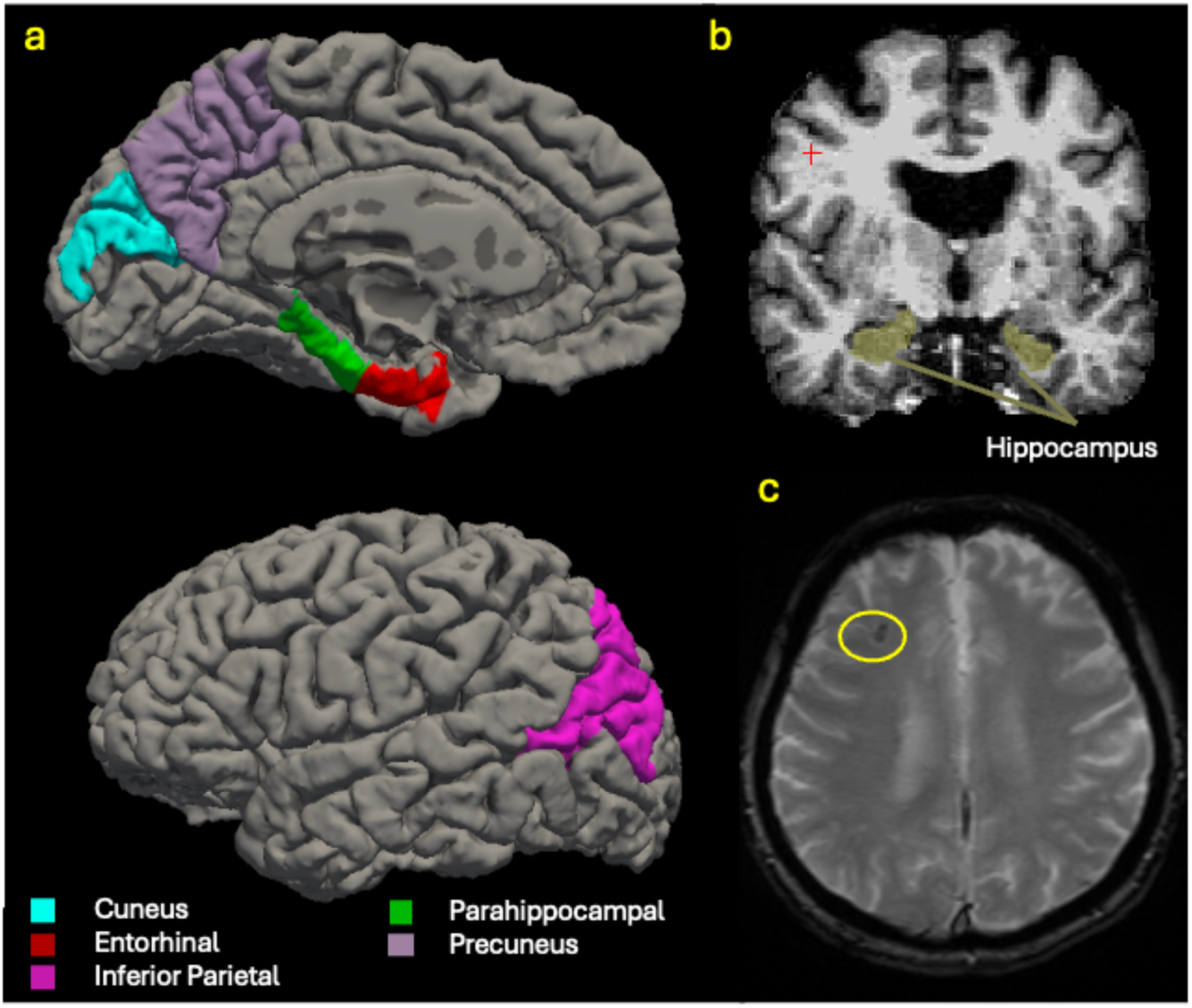
Anatomical features associated with Alzheimer’s disease. a. Alzheimer’s disease vulnerable regions mapped on the left hemisphere of a standard brain using a Freesurfer atlas b. Hippocampus, one of the six Alzheimer’s disease-vulnerable regions c. Cerebral microbleeds circled in yellow

Sleep deficiency, defined as having inappropriate quality, quantity, or timing of sleep, ^18^ is a robust risk factor associated with cognitive impairment and AD incidence. ^19–22^ Sleep architecture – the structural organization of sleep into rapid eye movement (REM) and non-REM sleep stages (N1, N2, N3 or slow wave sleep [SWS], and arousals) ^23^ – is an objective measure of sleep deficiency based on polysomnography, the gold standard for assessing sleep. While impaired sleep architecture (i.e., less SWS, less REM, and/or increased arousals) has been associated with AD diagnosis and cognitive decline, ^24,25^ little is known about the its association with the atrophy of AD-vulnerable regions and CMBs. Previously, exploratory whole-brain region of interest (ROI) analysis in young and middle-aged adults found less SWS and less non-REM sleep to be associated with lower gray matter thickness of some AD-vulnerable regions, including inferior-parietal lobule, precuneus, and cuneus. ^26,27^ In another study in middle-aged adults, lower density, amplitude, and duration of SWS and REM sleep were associated with smaller total gray matter volume. ^28^ However, these studies did not focus on AD-vulnerable regions or older adults, who have the highest risk of AD. Furthermore, none of the prior studies examined the association of sleep architecture with CMBs, although sleep apnea (i.e., a disorder where a person’s breathing stops multiple times during sleep and a cause of impaired sleep architecture) has been associated with increased risks of CMBs. ^29^ Finally, little is known about how sleep architecture synergizes with Apolipoprotein Ɛ4 (*APOE4*), the strongest known genetic risk factor of AD, ^30,31^ to affect the risk of brain atrophy and CMBs. Previously, *APOE4* carrying mice who were sleep deprived had higher levels of Aβ and tau pathologies than non-carrier sleep deprived mice, ^32^ suggesting that the neurological impact of impaired sleep architecture may be exacerbated in *APOE4* carriers.

This study examined the association of sleep architecture with brain atrophy in AD-vulnerable regions and the presence of CMBs in a sample with a high representation of older adults using the Atherosclerosis Risk in the Community Study (ARIC). In addition, we examined whether individuals with one or more *APOE4* alleles had increased brain atrophy and CMBs associated with impaired sleep architecture than non-carriers. Evidence generated by the current study will help characterize modifiable risk factors for AD and neurocognitive impairment by helping elucidate the contributions of sleep deficiency to distinct anatomical features that are associated with AD.

## METHODS

### Study Procedures and Analytic Sample

The Atherosclerosis Risk in the Communities Study (ARIC) is an ongoing investigation of health in community-living U.S. adults. Procedures for recruitment and data collection have been described in detail elsewhere. ^33^ Briefly, a subsample of ARIC participants took part in a sleep study between 1996-1998 as a part of the Sleep Heart Health Study, where they completed one night of unattended, in-home, overnight polysomnography. ^34^ Subsequently, a subset of surviving individuals who completed the polysomnography participated in a structural brain MRI protocol between 2011 and 2013. The protocols of ARIC have been approved by Institutional Review Boards (IRB) at all participating institutions: University of North Carolina at Chapel Hill IRB, Johns Hopkins University IRB, University of Minnesota IRB, and University of Mississippi Medical Center IRB.

The current study analyzed all ARIC participants who completed polysomnography and the structural brain MRI protocol. Exclusion criteria included having a stroke or probable dementia (defined as in prior ARIC studies by Mini Mental Status Examination score <21 if White and <19 if African-American). ^4^ Among 1,723 participants who had complete polysomnography data at baseline (i.e., the time when polysomnography was conducted), 294 participated in a substudy involving a structural MRI protocol. Of these individuals, two individuals were excluded due to previously having a stroke. None of the remaining participants had probable dementia. Subsequently, we excluded 14 participants without information on *APOE* genotype and 8 individuals with missing data on other covariates, making the final analytic sample size 270. Please see Supplementary Figure 1 for processes used to define the analytic sample. The current analyses have been approved by the Institutional Review Board at Yale University.

### Outcomes

Outcomes included (1) volumetric measurements of each AD-vulnerable region and (2) the presence of CMBs. Volumetric measurements were calculated based on images obtained using a T1-weighted magnetization-prepared rapid gradient-echo sequence (MPRAGE; 1.2-mm slices) on 3 Tesla Siemens scanners. ^4,35^ Detailed descriptions of MRI protocols are available from prior research. ^4,6,36–38^ Freesurfer was used to preprocess, reconstruct cortical surfaces, and conduct volumetric segmentation of the T1-weighted images. Each AD vulnerable region (i.e., hippocampus, parahippocampal, entorhinal, inferior parietal, precuneus and cuneus regions), selected based on prior evidence, which showed that atrophy specific to these regions was associated with an increased AD risk, ^2–6^ was defined based on the Freesurfer atlas. ^36,39,40^ The volume of each AD-vulnerable region equaled the sum of the respective region’s volume in the left and right hemispheres (in mm^3^). ^3–5^ These values were not corrected for intracranial volume.

Presence of CMBs was defined as having any hypointense regions in white or gray matter ≤10 mm in diameter. ^11,13^ CMBs were identified from images obtained using an axial T2* gradient recalled echo sequence (T2*GRE; 4-mm slices), also on 3 Tesla Siemens scanners. ^4,35^ All CMBs were identified by professional analysts and reviewed by a radiologist (interrater agreement=85%).^41^ For more details on the T2*GRE sequence and the identification of CMBs, please refer to prior research. ^4,6,36–38^ For the current research, we classified all participants dichotomously into those with at least one CMB and those without any CMBs. In addition, we examined the presence of lobar CMBs, which have been more closely associated with AD risk. ^15–17^ For this outcome, we classified individuals dichotomously into those with one or more lobar CMBs vs. those without any CMBs regardless of the type, excluding all individuals with any non-lobar [subcortical, deep cerebral, infratentorial] CMBs.

### Exposure

Compumedics PS polysomnography (Abbotsford, Victoria, Australia) was used to conduct full montage sleep monitoring according to published guidelines, based on comprehensive data obtained by electroencephalography (EEG), electrooculography, chin electromyography, thoracic and abdominal plethysmography, airflow, pulse oximetry, electrocardiography, and body position. ^34^ Using polysomnography data, sleep stages and arousals were manually scored on an epoch-by-epoch basis using standard criteria. ^34,42,43^ Please see prior research for detailed information on the sleep study’s procedures, data processing, and the scoring of sleep stages. ^34^ For the current study, exposures included the proportion of night-time sleep spent in SWS (henceforth referred to as SWS), the proportion of night-time sleep spent in REM sleep (henceforth referred to as REM), and arousal index (i.e., average number of arousals per hour) quantified from polysomnography. Proportions of time spent in SWS and REM were calculated by dividing time spent in the respective sleep stages with the total duration of night-time sleep. Arousal index was calculated by dividing the total number of arousals by total number of hours of night-time sleep (arousals per hour). Proportions of time spent in SWS and REM, rather than absolute durations of SWS or REM, were used to account for total sleep time.

### Covariates

Covariates included sex, age, years of education, apnea-hypopnea index (AHI; continuous), number of *APOE4* alleles, smoking status (never smoker, past smoker, current smoker), diabetes, body mass index, alcoholic drinks per week, hypertension, and baseline global cognitive performance. Global cognitive performance was quantified by taking an average of the standardized scores from the Delayed Word Recall, Digit Symbol Substitution, and Word Fluency tests, following prior research. ^44–46^ In addition to these baseline characteristics, we controlled for intracranial volume (estimated based on the T1 image using Freesurfer) ^47^ and whether a person had any heart failure events since the baseline polysomnography. We did not control for race-ethnicity as all participants in the analytic sample were White. Also, we did not adjust for depression as none of the participants had depression based on the Center for Epidemiologic Studies Depression Scale. ^48^

### Statistical analysis

Linear regression models examined the association of SWS, REM, and arousal index with the volume of each AD-vulnerable region, adjusting for all covariates. Separate models were used to examine each region as an outcome. We used false discovery rate to account for multiple comparisons as we investigated the volumes of six distinct brain regions as outcomes. The association of SWS, REM, and arousal index with the presence of CMBs was examined using logistic regression models. Separate models examined the presence of any CMB and that of lobar CMBs, respectively, as outcomes. If a measure of sleep architecture (i.e., SWS, REM, or arousal index) was significantly associated with an outcome, we additionally examined whether the association varied by *APOE* genotype using an interaction term. In addition, we examined the association of the following alternative measures of sleep with the anatomical features related to AD: absolute duration of slow wave sleep, absolute duration of rapid eye movement sleep, and wake after sleep onset (WASO), additionally accounting for time in bed.

For all analyses, statistical significance was determined based on a *p*-value of 0.05. All analysis included complete cases and were conducted using STATA (MP, version 17, College Station, TX).

## RESULTS

Table 1 shows sample characteristics, including demographics, AHI, *APOE* genotype, clinical characteristics, lifestyle choices, sleep architecture, and CMBs. Fifty-three percent were women, the median age was 61 (interquartile range [IQR]=57,66), all participants identified as White, and 47.0 % had 16 or more years of education. *APOE4* carriers (hetero- and homozygotes) accounted for 26.3%. The median proportions of time spent in SWS and REM sleep, respectively, were 17.4% and 21.5%. The median number of arousals was 17.1 per hour. Approximately one fourth of the participants had one or more CMBs. Individuals with lobar CMBs accounted for 8.2% of the full sample.

**Table 1.**
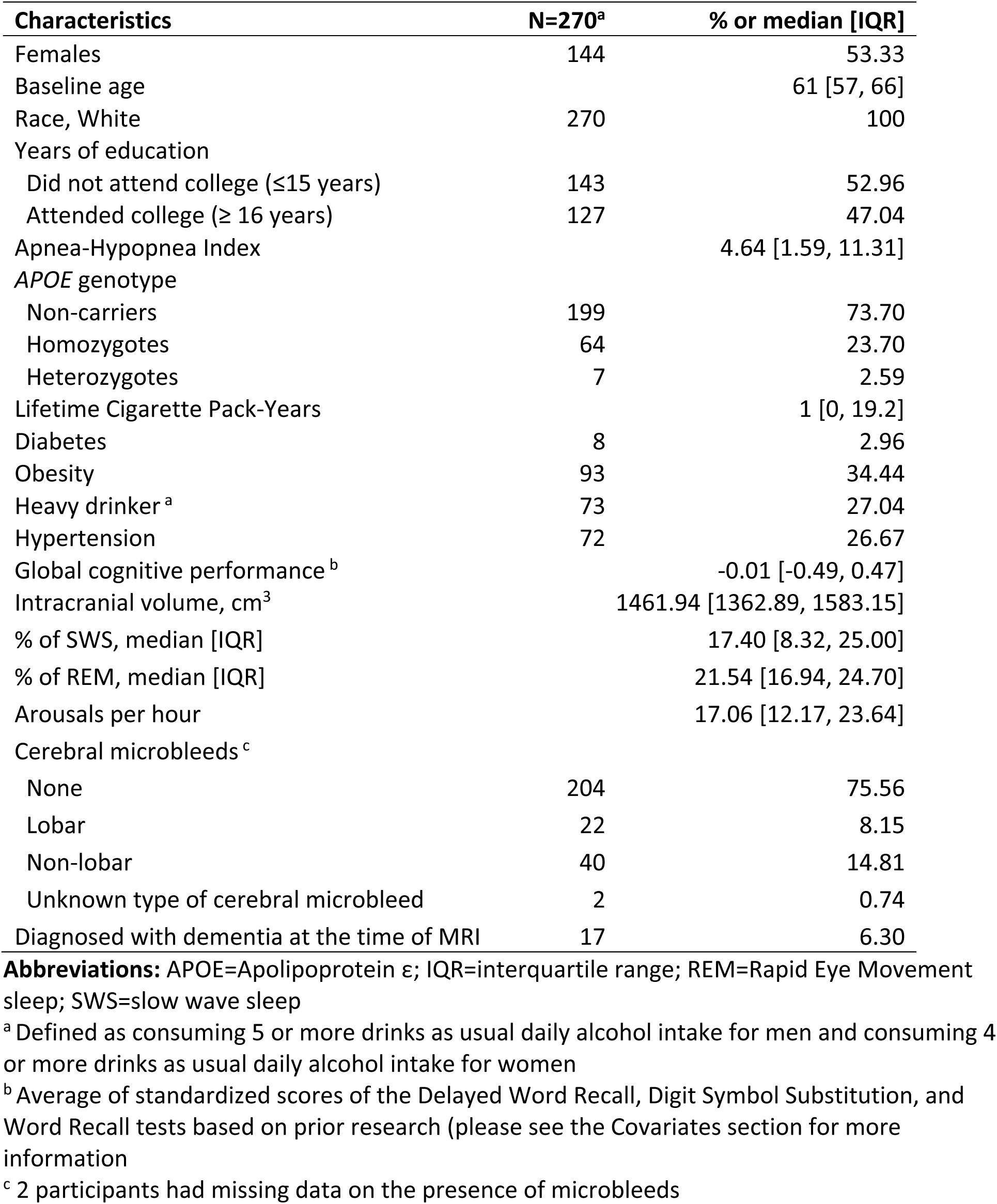
Sample characteristics.

Figure 2 depicts the association of sleep architecture (SWS, REM, arousal index) with the volume of each AD-vulnerable region, after adjusting for all covariates (see Supplementary Table 1 for minimally adjusted models). Having lower SWS was associated with smaller volumes of the inferior parietal region (β=−44.19 mm^3^ per percentage point [PP], 95%CI=−76.63, −11.76) and cuneus (β=−11.99 mm^3^ per PP, 95%CI=−20.93, −3.04). Having lower REM was associated with smaller volumes of the inferior parietal region (β=−75.52 mm^3^ per PP, 95%CI=−129.34, −21.70) and precuneus (β=−31.93 mm^3^ per PP, 95%CI=−63.79,−0.07). Having one more arousal per hour was associated with a slightly larger precuneus, β=25.62 mm^3^, 95%CI=0.85, 48.79. After an additional adjustment for multiple comparisons, inferior parietal volume remained significantly associated with both SWS and REM, respectively (false discovery rate-adjusted *p*s<0.05). The association between sleep architecture and inferior parietal volume did not vary by *APOE* genotype (Supplementary Table 2). Sensitivity analyses using alternative measures of sleep (i.e., absolute durations of SWS, REM, and WASO) showed comparable results, although statistical significance was attenuated (see Supplementary Table 3).

**Figure 2.**
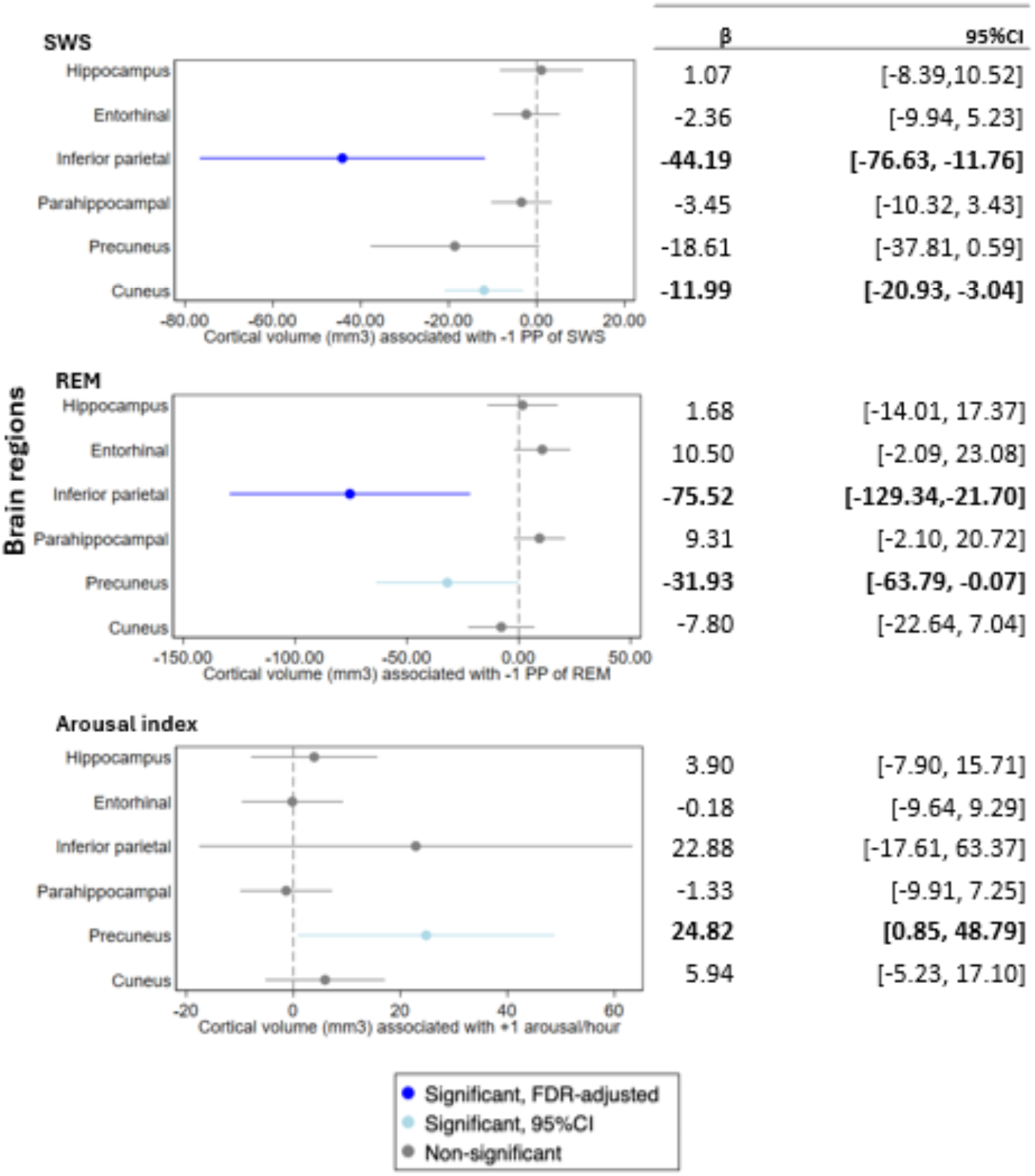
Cortical volume associated with sleep architecture for AD-vulnerable regions ^a^. **Abbreviations:** PP=percentage point; REM=Rapid Eye movement sleep; SWS=slow wave sleep Bold face indicates statistical significance based on 95% confidence intervals. * signifies statistical significance at the family-wise *p*<.05 after adjustments for false discovery rate. ^a^ All estimates are adjusted for sex, age, the number of *APOE4* alleles, years of education, apnea-hypopnea index, intracranial volume, global cognition, alcohol use, cigarette pack-years, hypertension, coronary heart disease (including myocardial infarction, heart failure, and percutaneous having done a revascularization procedure), diabetes, obesity, total sleep time, and multiple comparisons.

Table 2 shows the association of sleep architecture with the presence of CMBs, after adjustments for all covariates (see Supplementary Table 4 for unadjusted estimates). SWS, REM, and arousal index were not associated with the presence of any CMBs or that of lobar CMBs. Additional analyses using alternative measures of sleep (i.e., absolute durations of SWS and REM and WASO) yielded virtually identical estimates as the main analyses (Supplementary Table 5).

**Table 2.** Association between deficits in sleep architecture and the risk of cerebral microbleeds^a^.

| Outcomes | -1 PP of Slow wave sleep |  | -1 PP of REM sleep |  | +1/hour of Arousal index |  |
| --- | --- | --- | --- | --- | --- | --- |
|  | OR | 95%CI | OR | 95%CI | OR | 95%CI |
| Cerebral microbleeds<sup>b</sup> | 0.99 | [0.96, 1.03] | 1.01 | [0.96, 1.06] | 1.01 | [0.97, 1.05] |
| Lobar microbleeds<sup>c</sup> | 0.99 | [0.94, 1.05] | 1.04 | [0.96, 1.13] | 1.02 | [0.95, 1.09] |

**Abbreviations:** PP=percentage point; REM=Rapid Eye movement sleep
<sup>a</sup> All estimates are adjusted for sex, age, the number of *APOE4* alleles, years of education, apnea-hypopnea index, intracranial volume, global cognition, alcohol use, cigarette pack-years, hypertension, coronary heart disease (including myocardial infarction, heart failure, and percutaneous having done a revascularization procedure), diabetes, obesity, and total sleep time. Reference group for all models was 204 persons without CMBs.
<sup>b</sup> N=268. Two people were excluded due to missing data on the presence of CMBs.
<sup>c</sup> This model additionally excludes 40 individuals with non-lobar cerebral microbleeds, 2 individuals with missing data on cerebral microbleeds, and 5 individuals due to collinearity, resulting in N=221.

## DISCUSSION

In our sample of community-dwelling older adults, having a low proportion of SWS was associated with smaller inferior parietal and cuneus regions, even after adjusting for a large number of confounders. Having a lower proportion of REM sleep was also associated with smaller inferior parietal and precuneus regions. After additional adjustments for multiple comparisons, lower proportions of SWS and REM remained significantly associated with smaller volumes of the inferior parietal region. To the best of our knowledge, the current study is the first to examine the relationship of sleep architecture with the atrophy of AD-vulnerable regions in older adults. Previously, cortical thinning in parietal and temporal lobes has been observed during early, preclinical phases of AD, ^49,50^ but factors that precipitate the atrophy of these regions have remained obscure. Recent research has found shorter SWS and REM to be associated with lower total gray matter volume, ^28^ but did not examine distinct regions that had been shown to undergo increased neurodegeneration in preclinical and early AD. ^2–5^ Our research adds to this line of literature on sleep and AD by showing that lower SWS and REM may be precipitating factors of inferior parietal region atrophy, ^3,6^ although it should be noted that at least some of the observed atrophy may be unrelated to AD. Importantly, the current study’s findings can help characterize underlying mechanisms of how sleep deficiency, a prevalent disturbance among middle-aged and older adults, ^51–54^ may facilitate AD pathogenesis and cognitive impairment. Future studies need to investigate whether declines in cognitive domains associated with inferior parietal regions are associated with impairments in SWS and REM.

Furthermore, this research is the first to examine the association of sleep architecture with the presence of cerebral microbleeds. Contrary to our expectations, SWS, REM, arousals, and the alternative measures of sleep (i.e., absolute duration of SWS, absolute duration of REM, WASO) were not associated with the risk of CMBs or lobar CMBs. Our findings are at odds with prior research on sleep disordered breathing, which found sleep apnea (as defined by an AHI>=15), a known cause of impaired sleep architecture, to be associated with increased risks of CMBs. ^29^ Thus, it is possible that pathogenic features specific to sleep apnea that cannot sufficiently be captured by sleep architecture may significantly increase the risk of CMBs. ^55^ Previously, sleep apnea has been shown to be associated with cerebral desaturation, which can increase the risks of brain bleeding. ^56,57^ Also, sleep apnea is a risk factor for cerebral hypertension, which can lead to cerebral aneurysm enlargement, another risk factor for brain bleeding. ^58–60^ In comparison, impaired sleep architecture (i.e., shorter SWS and REM and higher arousals) is not directly related to cerebral desaturation or cerebral hypertension, and may have limited impact on brain bleeding independently from sleep disordered breathing. Further research in a sample enriched for CMBs is necessary as the rate of CMBs in our sample was relatively low, which may have reduced analytic power.

In addition, we examined whether the association of sleep architecture and the volumetric measurements of AD-vulnerable regions vary by *APOE* genotype. Previously, *APOE4* genotype was found to synergize with sleep disruption to facilitate Aβ accumulation and Aβ-associated tau seeding in mice. ^61^ A study of human adults also showed that not having consolidated sleep, defined as having increased number of arousals, was associated with higher risks of incident AD and tau pathology among *APOE4* carriers relative to non-carriers. ^22^ However, we did not find the association of sleep architecture with the volume of AD-vulnerable regions to vary by *APOE* genotype. Thus, *APOE4* carriers may not experience a heavier burden of brain atrophy associated with impaired sleep architecture. However, another possibility is that we did not find variations by *APOE4* because all individuals in our sample were White, which may introduce a bias as the participants may have had better access to preventive care for AD than underrepresented racial groups. ^62^ This possibility is supported by prior research that found Black *APOE4* carriers to experience heavier neurocognitive consequences of sleep apnea than Black non-carriers, ^63^ whereas among White adults, *APOE4* carriers experienced comparable neurocognitive consequences of sleep apnea as non-carriers. ^63^ Further research is warranted to examine racially diverse samples.

Finally, it remains unclear why cortical volume of the inferior parietal region was more strongly associated with SWS and REM than other AD-vulnerable brain regions. Prior research has shown that acute sleep deprivation alters metabolic activity in the inferior parietal region^64^ and cognitive performance requiring parietal activation. ^65^ Continued sleep deprivation may lead to impaired sleep architecture, which can, in turn, induce prolonged reduction in parietal activity, which has been associated with neurodegeneration. ^66^ It should also be noted that the association may be bidirectional, as cortical thinning of the parietal region has been associated with disturbances in SWS and REM. ^26^ Future research is needed to elucidate these associations.

The current study has several strengths. First, this study measured sleep deficiency using polysomnography, the gold-standard, and its association with the atrophy of regions that have been found to undergo early neurodegeneration in preclinical AD, thereby generating evidence on how sleep may be involved in AD pathogenesis. Second, this study is the first to examine the association of sleep architecture with cerebral microhemorrhages, an important predictor of stroke and dementia. Third, this study prospectively examined the association of baseline sleep architecture with brain atrophy and cerebral microbleeds, while controlling for baseline cognitive function. While this evidence is not causal, our findings provide preliminary evidence that short SWS and REM sleep may be observed in individuals prior to developing brain atrophy. Fourth, the current analyses adjusted for intracranial volume, *APOE* genotype, and a large number of demographic and clinical characteristics that are associated with both sleep deficiency and AD, thereby reducing the risks of confounding. Fifth, this study examined community-dwelling adults and its findings may be generalizable to non-institutionalized U.S. adults.

However, this study is not without its limitations. First, all individuals in our sample were White. Studies involving an assessment of overnight sleep architecture and MRI in underrepresented individuals are rare, despite evidence of their increased susceptibility to both AD and sleep disturbance. ^63,67^ Second, we used regional volumetric measures, but brain volume is associated with head size, which may have confounded our estimates. However, we adjusted our models for head size using total intracranial volume obtained from the structural MRI images. Third, as discussed previously, our analysis of CMBs may not have reached statistical significance due to limited power to analyze a dichotomous outcome, although this possibility is unlikely given that our point estimates are very close to the null. Fourth, our findings do not provide causal evidence, and research is necessary to examine whether changes in sleep deficiency may facilitate AD pathogenesis through contributing to the atrophy of the AD-vulnerable regions. To address these limitations, future research is needed to validate the current findings using a longitudinal study in a larger sample, which has a better representation of *APOE4* carriers and race-ethnically diverse populations.

## Conclusions

In a cohort of community-dwelling middle-aged and older adults, reduced SWS and REM, were associated with smaller inferior parietal region after adjusting for a large number of demographics, clinical characteristics, lifestyle choices, and multiple comparisons. Our findings provide preliminary evidence that reduced SWS and REM may contribute to brain atrophy, thereby increasing the risk of AD. Thus, sleep architecture may be a modifiable risk factor of AD and related dementias, warranting future research on underlying pathophysiological mechanisms.

## Abbreviations

APOE: Apolipoprotein ε
ARIC: Atherosclerosis Risk in the Communities Study
IQR: interquartile range
REM: Rapid Eye Movement sleep
SHHS: Sleep Heart Health Study
SWS: slow wave sleep

## Acknowledgments

This work was supported by the Claude D. Pepper Older Americans Independence Center at Yale’s Pilot Award (2P30AG021342-21). This Manuscript was prepared using ARIC and SHHS Research Materials obtained from the NHLBI Biologic Specimen and Data Repository Information Coordinating Center and does not necessarily reflect the opinions or views of the ARIC, SHHS or the NHLBI.

## Notes

None of the authors have financial support to declare and report absence of conflict of interest.

### Competing Interest Statement

The authors have declared no competing interest.

## REFERENCES

1. 2022 Alzheimer’s disease facts and figures. Alzheimers Dement. Apr 2022;18(4):700–789. doi:10.1002/alz.12638

2. Grothe MJ, Sepulcre J, Gonzalez-Escamilla G, et al. Molecular properties underlying regional vulnerability to Alzheimer’s disease pathology. Brain. 2018;141(9):2755–2771.

3. Dickerson BC, Bakkour A, Salat DH, et al. The Cortical Signature of Alzheimer’s Disease: Regionally Specific Cortical Thinning Relates to Symptom Severity in Very Mild to Mild AD Dementia and is Detectable in Asymptomatic Amyloid-Positive Individuals. Cerebral Cortex. 2008;19(3):497–510. doi:10.1093/cercor/bhn113

4. Knopman DS, Griswold ME, Lirette ST, et al. Vascular imaging abnormalities and cognition: mediation by cortical volume in nondemented individuals: atherosclerosis risk in communities-neurocognitive study. Stroke. Feb 2015;46(2):433–40. doi:10.1161/strokeaha.114.007847

5. Schwarz CG, Gunter JL, Wiste HJ, et al. A large-scale comparison of cortical thickness and volume methods for measuring Alzheimer’s disease severity. NeuroImage Clinical. 2016;11:802–812. doi:10.1016/j.nicl.2016.05.017

6. Dickerson BC, Stoub TR, Shah RC, et al. Alzheimer-signature MRI biomarker predicts AD dementia in cognitively normal adults. Neurology. Apr 19 2011;76(16):1395–402. doi:10.1212/WNL.0b013e3182166e96

7. Wu A, Sharrett AR, Gottesman RF, et al. Association of Brain Magnetic Resonance Imaging Signs With Cognitive Outcomes in Persons With Nonimpaired Cognition and Mild Cognitive Impairment. JAMA Netw Open. May 3 2019;2(5):e193359. doi:10.1001/jamanetworkopen.2019.3359

8. Karas G, Sluimer J, Goekoop R, et al. Amnestic mild cognitive impairment: structural MR imaging findings predictive of conversion to Alzheimer disease. Am J Neuroradiol. 2008;29(5):944–949.

9. Cai Z, Wang C, He W, et al. Cerebral small vessel disease and Alzheimer’s disease. Clin Interv Aging. 2015:1695–1704.

10. Kantarci K, Gunter JL, Tosakulwong N, et al. Focal hemosiderin deposits and β-amyloid load in the ADNI cohort. Alzheimers Dement. Oct 2013;9(5 Suppl):S116–23. doi:10.1016/j.jalz.2012.10.011

11. Akoudad S, Wolters FJ, Viswanathan A, et al. Association of Cerebral Microbleeds With Cognitive Decline and Dementia. JAMA neurology. Aug 1 2016;73(8):934–43. doi:10.1001/jamaneurol.2016.1017

12. Li X, Yuan J, Yang L, et al. The significant effects of cerebral microbleeds on cognitive dysfunction: an updated meta-analysis. PloS one. 2017;12(9):e0185145.

13. Martinez-Ramirez S, Greenberg SM, Viswanathan A. Cerebral microbleeds: overview and implications in cognitive impairment. Alzheimers Res Ther. 2014/06/11 2014;6(3):33. doi:10.1186/alzrt263

14. Ni J, Auriel E, Martinez-Ramirez S, et al. Cortical localization of microbleeds in cerebral amyloid angiopathy: an ultra high-field 7T MRI study. J Alzheimers Dis. 2015;43(4):1325–30. doi:10.3233/jad-140864

15. Yates PA, Villemagne VL, Ellis KA, Desmond PM, Masters CL, Rowe CC. Cerebral microbleeds: a review of clinical, genetic, and neuroimaging associations. Front Neurol. Jan 6 2014;4:205. doi:10.3389/fneur.2013.00205

16. Yates PA, Sirisriro R, Villemagne VL, Farquharson S, Masters CL, Rowe CC. Cerebral microhemorrhage and brain β-amyloid in aging and Alzheimer disease. Neurology. Jul 5 2011;77(1):48–54. doi:10.1212/WNL.0b013e318221ad36

17. Yates PA, Desmond PM, Phal PM, et al. Incidence of cerebral microbleeds in preclinical Alzheimer disease. Neurology. 2014;82(14):1266–1273.

18. What Are Sleep Deprivation and Deficiency? National Heart, Lung, and Blood Institute. https://www.nhlbi.nih.gov/health/sleep-deprivation#:~:text=Sleep%20deficiency%20can%20interfere%20with,or%20worried%20in%20social%20situations.

19. Blackman J, Love S, Sinclair L, Cain R, Coulthard E. APOE ε4, Alzheimer’s disease neuropathology and sleep disturbance, in individuals with and without dementia. Alzheimers Res Ther. Mar 30 2022;14(1):47. doi:10.1186/s13195-022-00992-y

20. Cavaillès C, Carrière I, Wagner M, et al. Trajectories of sleep duration and timing before dementia: a 14-year follow-up study. Age Ageing. Aug 2 2022;51(8)doi:10.1093/ageing/afac186

21. Cipriani G, Lucetti C, Danti S, Nuti A. Sleep disturbances and dementia. Psychogeriatrics. Mar 2015;15(1):65–74. doi:10.1111/psyg.12069

22. Lim AS, Kowgier M, Yu L, Buchman AS, Bennett DA. Sleep fragmentation and the risk of incident Alzheimer’s disease and cognitive decline in older persons. Sleep. 2013;36(7):1027–1032.

23. Practice parameters for the indications for polysomnography and related procedures. Polysomnography Task Force, American Sleep Disorders Association Standards of Practice Committee. Sleep. Jun 1997;20(6):406–22.

24. Himali JJ, Baril A-A, Cavuoto MG, et al. Association Between Slow-Wave Sleep Loss and Incident Dementia. JAMA neurology. 2023;doi:10.1001/jamaneurol.2023.3889

25. Pase MP, Himali JJ, Grima NA, et al. Sleep architecture and the risk of incident dementia in the community. Neurology. Sep 19 2017;89(12):1244–1250. doi:10.1212/wnl.0000000000004373

26. Dubé J, Lafortune M, Bedetti C, et al. Cortical thinning explains changes in sleep slow waves during adulthood. J Neurosci. May 20 2015;35(20):7795–807. doi:10.1523/jneurosci.3956-14.2015

27. Weihs A, Frenzel S, Garvert L, et al. The relationship between Alzheimer’s-related brain atrophy patterns and sleep macro-architecture. Alzheimers Dement. 2022;14(1):e12371.

28. Baril A-A, Beiser AS, Mysliwiec V, et al. Slow-Wave Sleep and MRI Markers of Brain Aging in a Community-Based Sample. Neurology. 2021;96(10):e1462–e1469. doi:10.1212/wnl.0000000000011377

29. Koo DL, Kim JY, Lim JS, Kwon HM, Nam H. Cerebral Microbleeds on MRI in Patients with Obstructive Sleep Apnea. J Clin Sleep Med. Jan 15 2017;13(1):65–72. doi:10.5664/jcsm.6390

30. Elias-Sonnenschein LS, Viechtbauer W, Ramakers IH, Verhey FR, Visser PJ. Predictive value of APOE-ε4 allele for progression from MCI to AD-type dementia: a meta-analysis. J Neurol Neurosurg Psychiatry. Oct 2011;82(10):1149–56. doi:10.1136/jnnp.2010.231555

31. Gharbi-Meliani A, Dugravot A, Sabia S, et al. The association of APOE ε4 with cognitive function over the adult life course and incidence of dementia: 20 years follow-up of the Whitehall II study. Alzheimers Res Ther. Jan 4 2021;13(1):5. doi:10.1186/s13195-020-00740-0

32. Wang C, Nambiar A, Strickland MR, et al. APOE-ε4 synergizes with sleep disruption to accelerate Aβ deposition and Aβ-associated tau seeding and spreading. J Clin Invest. Jul 17 2023;133(14)doi:10.1172/jci169131

33. Investigators. The Atherosclerosis Risk in Communities (ARIC) Study: design and objectives. The ARIC investigators. Am J Epidemiol. Apr 1989;129(4):687–702.

34. Quan SF, Howard BV, Iber C, et al. The Sleep Heart Health Study: design, rationale, and methods. Sleep. Dec 1997;20(12):1077–85.

35. Moazzami K, Shao IY, Chen LY, et al. Atrial Fibrillation, Brain Volumes, and Subclinical Cerebrovascular Disease (from the Atherosclerosis Risk in Communities Neurocognitive Study [ARIC-NCS]). Am J Cardiol. Jan 15 2020;125(2):222–228. doi:10.1016/j.amjcard.2019.10.010

36. Desikan RS, Ségonne F, Fischl B, et al. An automated labeling system for subdividing the human cerebral cortex on MRI scans into gyral based regions of interest. NeuroImage. Jul 1 2006;31(3):968–80. doi:10.1016/j.neuroimage.2006.01.021

37. Jones DT, Machulda MM, Vemuri P, et al. Age-related changes in the default mode network are more advanced in Alzheimer disease. Neurology. Oct 18 2011;77(16):1524–31. doi:10.1212/WNL.0b013e318233b33d

38. ARIC Analysis Manual 30 (Version 3.0). https://aric.cscc.unc.edu/aric9/sites/default/files/public/visitdocuments/v9/ARIC_Manual_30.pdf

39. Fischl B. FreeSurfer. NeuroImage. Aug 15 2012;62(2):774–81. doi:10.1016/j.neuroimage.2012.01.021

40. Fischl B, Salat DH, Busa E, et al. Whole brain segmentation: automated labeling of neuroanatomical structures in the human brain. Neuron. Jan 31 2002;33(3):341–55. doi:10.1016/s0896-6273(02)00569-x

41. Kantarci K, Gunter JL, Tosakulwong N, et al. Focal hemosiderin deposits and β-amyloid load in the ADNI cohort. Alzheimers Dement. 2013;9(5):S116–S123.

42. EEG arousals: scoring rules and examples: a preliminary report from the Sleep Disorders Atlas Task Force of the American Sleep Disorders Association. Sleep. Apr 1992;15(2):173–84.

43. A manual of standardized terminology, techniques and scoring system for sleep stages of human subjects. Allan Rechtschaffen and Anthony Kales, editors. In: Kales A, Rechtschaffen A, editors. Bethesda, Md.: U. S. National Institute of Neurological Diseases and Blindness, Neurological Information Network; 1968.

44. Deal JA, Sharrett AR, Albert MS, et al. Hearing Impairment and Cognitive Decline: A Pilot Study Conducted Within the Atherosclerosis Risk in Communities Neurocognitive Study. Am J Epidemiol. 2015;181(9):680–690. doi:10.1093/aje/kwu333

45. Gottesman RF, Rawlings AM, Sharrett AR, et al. Impact of differential attrition on the association of education with cognitive change over 20 years of follow-up: the ARIC neurocognitive study. Am J Epidemiol. 2014;179(8):956–966.

46. Gottesman RF, Schneider AL, Albert M, et al. Midlife hypertension and 20-year cognitive change: the atherosclerosis risk in communities neurocognitive study. JAMA neurology. 2014;71(10):1218–1227.

47. Buckner RL, Head D, Parker J, et al. A unified approach for morphometric and functional data analysis in young, old, and demented adults using automated atlas-based head size normalization: reliability and validation against manual measurement of total intracranial volume. NeuroImage. 2004;23(2):724–738.

48. Lewinsohn PM, Seeley JR, Roberts RE, Allen NB. Center for Epidemiologic Studies Depression Scale (CES-D) as a screening instrument for depression among community-residing older adults. Psychol Aging. 1997;12(2):277.

49. Jacobs HI, Van Boxtel MP, Jolles J, Verhey FR, Uylings HB. Parietal cortex matters in Alzheimer’s disease: an overview of structural, functional and metabolic findings. Neurosci Biobehav Rev. Jan 2012;36(1):297–309. doi:10.1016/j.neubiorev.2011.06.009

50. Greene SJ, Killiany RJ. Subregions of the inferior parietal lobule are affected in the progression to Alzheimer’s disease. Neurobiol Aging. Aug 2010;31(8):1304–11. doi:10.1016/j.neurobiolaging.2010.04.026

51. Miner B, Doyle M, Knauert M, et al. Insomnia with objective short sleep duration in community-living older persons: A multifactorial geriatric health condition. J Am Geriatr Soc. Apr 2023;71(4):1198–1208. doi:10.1111/jgs.18195

52. Miner B, Gill TM, Yaggi HK, et al. Insomnia in Community-Living Persons with Advanced Age. J Am Geriatr Soc. Aug 2018;66(8):1592–1597. doi:10.1111/jgs.15414

53. Miner B, Gill TM, Yaggi HK, et al. The Epidemiology of Patient-Reported Hypersomnia in Persons With Advanced Age. J Am Geriatr Soc. Dec 2019;67(12):2545–2552. doi:10.1111/jgs.16107

54. Miner B, Kryger MH. Sleep in the aging population. Sleep Med Clin. 2020;15(2):311–318.

55. Aktas O, Ullrich O, Infante-Duarte C, Nitsch R, Zipp F. Neuronal damage in brain inflammation. Arch Neurol. 2007;64(2):185–189.

56. Burzyńska M, Uryga A, Kasprowicz M, Czosnyka M, Dragan B, Kübler A. The relationship between the time of cerebral desaturation episodes and outcome in aneurysmal subarachnoid haemorrhage: a preliminary study. J Clin Monit Comput. Aug 2020;34(4):705–714. doi:10.1007/s10877-019-00377-x

57. Zhang Z, Qi M, Hügli G, Khatami R. Predictors of changes in cerebral perfusion and oxygenation during obstructive sleep apnea. Sci Rep. 2021/12/06 2021;11(1):23510. doi:10.1038/s41598-021-02829-4

58. Kohler M, Pepperell J, Casadei B, et al. CPAP and measures of cardiovascular risk in males with OSAS. Eur Resp J. 2008;32(6):1488–1496.

59. Pepperell JC, Ramdassingh-Dow S, Crosthwaite N, et al. Ambulatory blood pressure after therapeutic and subtherapeutic nasal continuous positive airway pressure for obstructive sleep apnoea: a randomised parallel trial. The Lancet. 2002;359(9302):204–210.

60. Zaremba S, Albus L, Schuss P, Vatter H, Klockgether T, Güresir E. Increased risk for subarachnoid hemorrhage in patients with sleep apnea. J Neurol. 2019;266:1351–1357.

61. Wang C, Nambiar A, Strickland MR, et al. APOE-ε4 synergizes with sleep disruption to accelerate Aβ deposition and Aβ-associated tau seeding and spreading. J Clin Investig. 2023;

62. Minorities and Women Are at Greater Risk for Alzheimer’s Disease. https://www.cdc.gov/aging/publications/features/Alz-Greater-Risk.html.

63. Turner AD, Locklear CE, Oruru D, Briggs AQ, Bubu OM, Seixas A. Exploring the combined effects of sleep apnea and APOE-e4 on biomarkers of Alzheimer’s disease. Front Aging Neurosci. 2022;14:1017521. doi:10.3389/fnagi.2022.1017521

64. Thomas M, Sing H, Belenky G, et al. Neural basis of alertness and cognitive performance impairments during sleepiness. I. Effects of 24 h of sleep deprivation on waking human regional brain activity. J Sleep Res. Dec 2000;9(4):335–52. doi:10.1046/j.1365-2869.2000.00225.x

65. Ma N, Dinges DF, Basner M, Rao H. How acute total sleep loss affects the attending brain: a meta-analysis of neuroimaging studies. Sleep. 2015;38(2):233–240.

66. Zilberter Y, Zilberter M. The vicious circle of hypometabolism in neurodegenerative diseases: ways and mechanisms of metabolic correction. J Neurosci Res. 2017;95(11):2217–2235.

67. Ahn S, Lobo JM, Logan JG, Kang H, Kwon Y, Sohn MW. A scoping review of racial/ethnic disparities in sleep. Sleep Med. May 2021;81:169–179. doi:10.1016/j.sleep.2021.02.027

