## Supplementary files for "Lower slow wave sleep and rapid eye-movement sleep are associated with brain atrophy of AD-vulnerable regions"

**Supplementary Materials**

**Figure S1.** Identification of the analytic sample

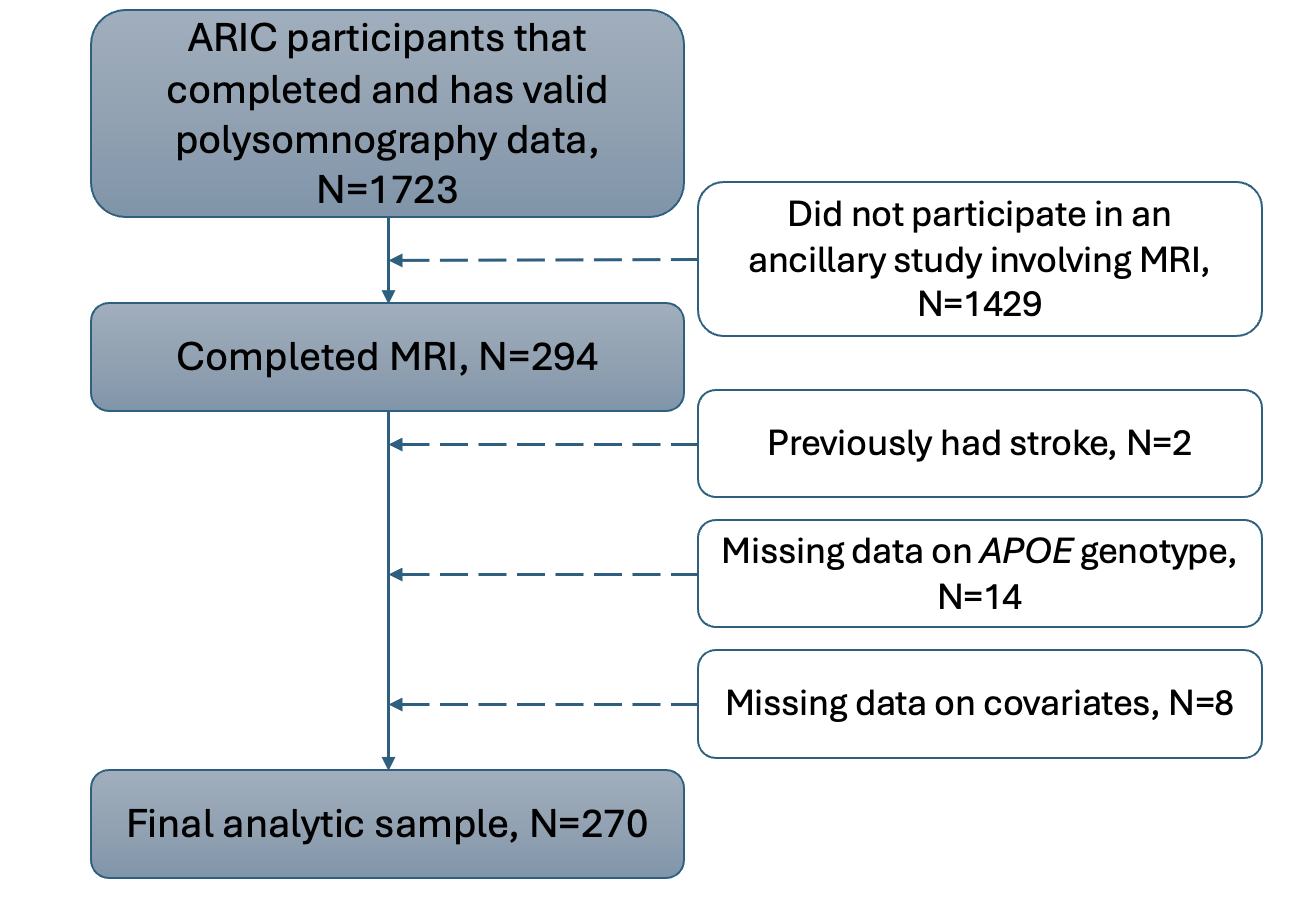

**Abbreviations:** ARIC=Atherosclerosis Risk in the Community Study

**Table S1.** Minimally adjusted association between impaired sleep architecture and brain volumes of AD-vulnerable regions ^a^

| **Region** | **Cortical volume associated with 1 unit of sleep index** | | | | | | | | | | | | | | | | | | | | | |
| --- | --- | --- | --- | --- | --- | --- | --- | --- | --- | --- | --- | --- | --- | --- | --- | --- | --- | --- | --- | --- | --- | --- |
|  | **-1 PP of Slow wave sleep** | | | | | |  | **-1 PP of REM sleep** | | | | | | |  | | **+1/hour of Arousal index** | | | | | |
|  | **β** | **95%CI** | | | | |  | **β** | **95%CI** | | | | | |  | | **β** | **95%CI** | | | | |
| Hippocampus | 5.33 | [ | -5.40 | , | 16.06 | ] |  | -1.49 | [ | -19.82 | , | 16.83 | ] |  | | -0.87 | | [ | -11.81 | , | 10.06 | ] |
| Entorhinal | 2.06 | [ | -5.66 | , | 9.78 | ] |  | 9.06 | [ | -4.12 | , | 22.25 | ] |  | | -1.17 | | [ | -9.04 | , | 6.69 | ] |
| Inferior Parietal | **-33.62** | **[** | **-66.03** | **,** | **-1.20** | **]** |  | **-79.90** | **[** | **-135.27** | **,** | **-24.52** | **]** |  | | 18.59 | | [ | -14.45 | , | 51.63 | ] |
| Parahippocampal | -2.01 | [ | -8.97 | , | 4.96 | ] |  | 6.17 | [ | -5.73 | , | 18.07 | ] |  | | -2.05 | | [ | -9.15 | , | 5.05 | ] |
| Precuneus | -14.62 | [ | -34.08 | , | 4.83 | ] |  | **-34.23** | **[** | **-67.47** | **,** | **-0.99** | **]** |  | | 15.60 | | [ | -4.23 | , | 35.44 | ] |
| Cuneus | **-9.04** | **[** | **-17.60** | **,** | **-0.48** | **]** |  | -12.46 | [ | -27.09 | , | 2.17 | ] |  | | 6.43 | | [ | -2.30 | , | 15.16 | ] |

**Abbreviations:** PP=percentage point; REM=Rapid Eye movement sleep

Bold face indicates statistical significance based on 95% confidence intervals.

* (asterisk) signifies statistical significance at the level of *p*<.05

^a^ These estimates are adjusted only for total intracranial volume. N=269

**Table S2.** Variations in the associations of deficits in sleep architecture and inferior parietal region volume by *APOE* genotype

| **Estimate** | **-1 PP of slow wave sleep** | | | | | |  | | **-1 PP of REM sleep** | | | | | | | |
| --- | --- | --- | --- | --- | --- | --- | --- | --- | --- | --- | --- | --- | --- | --- | --- | --- |
|  | **β** | **95%CI** | | | | |  | | **β** | | **95%CI** | | | | | |
| Inferior parietal region volume | **-108.89** | **[** | **-188.79** | **,** | **-28.99** | **]** | |  | | -79.37 | | [ | -222.92 | , | 64.18 | ] |
| Difference associated with having 1 more APOE4 allele | 51.75 | [ | -6.71 | , | 110.21 | ] | |  | | 3.04 | | [ | -101.99 | , | 108.06 | ] |

**Abbreviations:** PP=percentage point; REM=Rapid Eye movement sleep

Bold face denotes statistical significance at the level of 0.05.

^a^ Indicates reductions in cortical volume associated with 1 unit decrease in the respective sleep index

**Table S3.** Associations of deficits in alternative measures of sleep with cortical volume of AD vulnerable regions

| **Brain region** | **-1 min of SWS duration** | | | | | | |  | **-1 min of REM duration** | | | | | | |  | **+1 min of WASO** | | | | | |
| --- | --- | --- | --- | --- | --- | --- | --- | --- | --- | --- | --- | --- | --- | --- | --- | --- | --- | --- | --- | --- | --- | --- |
|  | **β** |  | **95%CI** | | | | |  | **β** |  | **95%CI** | | | | |  | **β** | **95%CI** | | | | |
| Hippocampus | 0.58 |  | [ | -1.87 | , | 3.02 | ] |  | 1.11 |  | [ | -3.07 | , | 5.29 | ] |  | 0.67 | [ | -2.01 | , | 3.35 | ] |
| Entorhinal | -0.53 |  | [ | -2.49 | , | 1.43 | ] |  | **3.50** |  | **[** | **0.16** | **,** | **6.84** | **]** |  | -0.87 | [ | -3.01 | , | 1.28 | ] |
| Inferior Parietal | **-9.54** |  | **[** | **-17.96** | **,** | **-1.11** | **]** |  | **-19.26** |  | **[** | **-33.64** | **,** | **-4.88** | **]** |  | 0.18 | [ | -9.06 | , | 9.42 | ] |
| Parahippocampal | -0.67 |  | [ | -2.44 | , | 1.10 | ] |  | 2.53 |  | [ | -0.50 | , | 5.55 | ] |  | -1.74 | [ | -3.68 | , | 0.20 | ] |
| Precuneus | -3.63 |  | [ | -8.63 | , | 1.37 | ] |  | -6.84 |  | [ | -15.37 | , | 1.69 | ] |  | 3.00 | [ | -2.48 | , | 8.48 | ] |
| Cuneus | -2.29 |  | [ | -4.63 | , | 0.04 | ] |  | -1.37 |  | [ | -5.34 | , | 2.61 | ] |  | 0.20 | [ | -2.36 | , | 2.75 | ] |

**Abbreviations:** SWS=slow wave sleep; REM=rapid eye movement sleep; WAS=wake after sleep onset

Bold face indicates statistical significance based on 95% confidence intervals.

**Table S4.** Unadjusted association between deficits in sleep architecture and the risk of cerebral microbleeds

| **Outcomes** | **-1PP of Slow wave sleep** | | |  | **-1PP of REM sleep** | |  | **+1/hour of Arousal index** | |
| --- | --- | --- | --- | --- | --- | --- | --- | --- | --- |
|  | **OR** |  | **95%CI** |  | **OR** | **95%CI** |  | **OR** | **95%CI** |
| Any Cerebral microbleeds ^a^ | 1.01 |  | [0.98, 1.03] |  | 1.01 | [0.96,1.06] |  | 1.00 | [0.98, 1.03] |
| Lobar microbleeds ^b^ | 1.02 |  | [0.97, 1.06] |  | 1.03 | [0.95, 1.11] |  | 0.99 | [0.94, 1.04] |

**Abbreviations:** REM=Rapid Eye movement sleep

Bold face indicates statistical significance based on 95% confidence intervals.

^a^ N=268 after excluding 2 individuals with missing data on cerebral microbleeds.

^b^ This model excludes 40 individuals with non-lobar CMBs, 2 individuals with missing data on cerebral microbleeds, and 5 individuals due to collinearity, resulting in N=221.

**Table S5.** Association of deficits in the alternative measures of sleep and the risk of cerebral microbleeds

| **Cerebral microbleeds** | **-1 min of SWS duration** | | |  | **-1 min of REM duration** | | |  | **+1 min of WASO** | |
| --- | --- | --- | --- | --- | --- | --- | --- | --- | --- | --- |
|  | **β** |  | **95%CI** |  | **β** |  | **95%CI** |  | **β** | **95%CI** |
| Any cerebral microbleeds ^a^ | 1.00 |  | [0.99,1.01] |  | 1.00 |  | [0.99,1.02] |  | 1.00 | [0.99,1.01] |
| Lobar microbleeds ^b^ | 1.00 |  | [0.99,1.01] |  | 1.02 |  | [0.99,1.04] |  | 0.99 | [0.97,1.01] |

**Abbreviations:** min=minute; REM=Rapid Eye movement sleep; SWS=slow wave sleep; WASO=wake after sleep onset

^a^ N=269 after excluding 2 individuals with missing data on cerebral microbleeds.

^b^ This model excludes 40 individuals with non-lobar CMBs, 2 individuals with missing data on cerebral microbleeds, and 5 individuals due to collinearity, resulting in N=219.
